# Perceptions and prospects in life sciences in a heterogeneous Latin American population

**DOI:** 10.1101/514216

**Authors:** Leonardo M.R. Ferreira, Giovanni A. Carosso, Bruno Lopez-Videla, Gustavo Vaca Diez, Laura Ines Rivera-Betancourt, Yara Rodriguez, Dalila G. Ordonez, Natalia Montellano Duran, Diana K. Alatriste-Gonzalez, Aldo Vacaflores, Soad Bohorquez, Lilian Gonzalez Auza, Christian Schuetz, Carolina V. Alexander-Savino, Omar Gandarilla Cuellar, Mohammed A. Mostajo-Radji

**Affiliations:** Clubes de Ciencia Bolivia Foundation, Santa Cruz de la Sierra, Bolivia.; Department of Surgery and Diabetes Center, University of California San Francisco, San Francisco, CA, USA; McKusick-Nathans Institute of Genetic Medicine, Johns Hopkins University School of Medicine, Baltimore, MD, USA; Clinical Sequencing Laboratory, Italian Hospital of Buenos Aires, Buenos Aires, Argentina; Department of Personality, Evaluation and Psychology Treatments, Psychology, Salamanca University, Salamanca, Spain; Feinberg School of Medicine, Northwestern University, Chicago, IL, USA; Department of Molecular and Cellular Biology, Harvard University, Cambridge, MA, USA; Institute for Cell and Neurobiology, Charité CrossOver, Charité Universitätsmedizin Berlin, Berlin, Germany; Department of Cell Biology, Harvard Medical School, Boston, MA, USA; Institute of Medical Biotechnology, Technological University of Berlin, Berlin, Germany; Center for Transplantation Sciences, Massachusetts General Hospital, Boston, MA, USA; Rochester General Hospital Research Institute, Rochester, NY, USA; Beth Israel Deaconess Medical Center, Boston, MA, USA

**Keywords:** Science Education, Biology Education, Bolivia, Developing Countries, Latin America

## Abstract

Particular challenges exist for science education in the developing world, where limited resources beget curricula designed to balance state-of-the-art knowledge with practical and political considerations in region-specific contexts. Project-based biology teaching is particularly difficult to execute due to high infrastructural costs and limited teacher training. Here, we report our results implementing short, challenging, and low-cost biology courses to high school and college students in Bolivia, designed and taught in collaboration between scientists from developed nations and local science instructors. We find our approach to be effective at transmitting advanced topics in disease modeling, microscopy, genome engineering, neuroscience, microbiology, and regenerative biology. Importantly, this approach was unaffected by the students’ backgrounds, education level, socioeconomic status, or initial interest in the course, and increased participants’ interest in pursuing scientific careers. These results demonstrate efficacy of participatory learning in a developing nation, and suggest that such techniques could drive scientific engagement in other developing economies.

## INTRODUCTION

Latin America is home to more than 10% of the world’s population, but accounts for only 5% of the global scientific output [1-4], as measured by quantity of original research publications and patent awards. A key indicator of performance in science, technology, and innovation (STI), these output metrics suggest significant underrepresentation of the region on the global stage. Even outside of Latin America, Latinos tend to be underrepresented in scientific careers. For example, in the United States, while Latinos comprise over 17% of the country’s population, they receive under 6% of graduate degrees in science, technology, engineering, and mathematics (STEM) fields [5]. Importantly, while Latinos are among the fastest growing demographics in the developed world, their participation in higher education, and science in particular, is not projected to keep pace with this growth [6]. Several intervention strategies have been deployed in an attempt to minimize these gaps, but disproportionately low Latino representation in STEM remains a conundrum for policymakers, universities, and science educators.

Competitive STI performance is a critical determinant of sustainable economic development. Within Latin America, the Plurinational State of Bolivia represents a particularly interesting case for examining individual STI engagement. Bolivia’s 36 indigenous groups comprise over 70% of the total population, making this relatively small territory the most diverse country in Latin America [7, 8]. Moreover, three Bolivian cities are projected to be the fastest-growing Latin American economies by 2030, as measured by middle-income population growth, with a fifteen-fold hike expected in the most populous city, Santa Cruz [9]. However, the country accounts for only 0.22% of Latin American STI research outputs and has negligible rates of patent awards [4, 10]. Despite a decade of relative political stability and government education expenditures exceeding 8% of GDP—highest in the region and far above any developed country, with the United States, United Kingdom, and France investing less than 6% of GDP [11-13] — Bolivia consistently places last in the region according to World Intellectual Property Organization (WIPO) and Global Innovation Index measures [14, 15]. The country does not implement standardized education metrics such as the Program for International Student Assessment (PISA), further obfuscating causal analyses and international oversight [16] and raising a need for alternative methods of analysis.

Several efforts have been carried out in Bolivia to increase student retention and interest in education. K-12 education in Bolivia is, in principle, mandatory for all Bolivian citizens, and the public system is completely free for all students. Since 2006, the Bolivian government has implemented a nation-wide conditional cash transfer program, locally known as “Bono Juancito Pinto”, to incentivize continuing education [17]. Public university enrollment requires only a nominal student fee (equivalent to less than $30 per semester), and its funding is guaranteed as a percentage of revenue from nationalized oil and gas sales. Public universities have complete administrative autonomy from the central government. Yet, despite secured funding and freedom to appoint their governing body and faculty, Bolivian universities perform poorly in world rankings and experience low student retention rates [18]. Indeed, Bolivia is the only Latin American nation with dropout rates in excess of completion rates for higher education [18].

The Bolivian case is illustrative of a central challenge in science education in developing countries, where resource limitations and outsize political influence in curriculum development often result in compromised schooling priorities, which manifests in three distinct ways. First, resource limitations precipitate a compromise between instruction of curricula with direct practicality in the local context and curricula reflecting the most recent or cutting-edge findings in developing nations [19, 20]. Cell biology, for instance, in Bolivia may take a backseat to more tangible lessons in Incan medicinal traditions. Second, lessons with relevance to particular political agendas may receive priority over more foundational subject matter. The 13-pronged national development directive, for example, contains an emphasis on “wisdom of the ancients,” while mentioning “science and technology” only in a call for national sovereignty therein. Moreover, the 2010 education reform “Avelino Siñani - Elizardo Perez” highlighted three conceptual elements of decolonization, community education, and “productive education”, while disregarding teaching methodology and training of teachers [21, 22]. Lastly, pressures to cover particular topics, and quickly progress to the next, often result in superficial student understanding on the basis of rote, memorization-based knowledge. In other words, science education policies largely focus on what to teach, but neglect how to teach [23]. Thus, a typical Bolivian student is incentivized toward simple repetition of facts, but discouraged from asking questions or establishing discussions with faculty [21, 24, 25].

Project-based teaching has been shown to be particularly effective at transmitting scientific concepts [26, 27], especially to students from underrepresented communities [28, 29]. In this approach, students are given complex tasks to be resolved over a defined period of time and are expected to conclude with a finalized product or a presentation [30]. Educational outcome data suggest that such “hands-on” approaches to teaching would be particularly adapted to the needs of students in developing nations, yet their implementation has been limited [31], with the majority of studies on the impact of project-based teaching in underrepresented communities having been performed in the developed world [29, 32]. Today, comparatively few studies have been carried out to assess the potential educational impact of project-based course adoption in Latin America in a region-specific manner.

Here, we sought to determine whether a grassroots approach, which we named Clubes de Ciencia Bolivia (CdeCBo), centered around participatory learning could impact individual students’ motivation toward STEM careers and at the same time provide feedback regarding perceptions thereof, among Bolivian youth. By exploiting national and social media outlets, we were able to reach over 50% of Bolivian youth with internet access. Notably, we found that, despite high baseline motivation and confidence, as well as a vastly positive perception of education quality, students performed poorly when tested on basic knowledge in biology. We further show that, upon completion of hands-on, hybrid lecture and project-based workshops and exposure to the most recent scientific developments, students demonstrated rapid’ learning of new knowledge, as well as high scores in confidence and enthusiasm in relation to their own prospects in pursuit of STEM careers. Our results indicate that grassroots-level workshops, fostering personalized collaboration between highly-trained instructors and local students, are an effective means of driving STEM motivation across a diverse youth population, and suggest that such platforms may prove fruitful in a diverse range of developing economies.

## METHODS

### Survey design

A survey was designed to interrogate the experience and perceptions in science education of Bolivian high school and college students nationwide. The survey contained 5 sections. The first set of questions pertained to basic demographic information, such as department of residence, grade in which they were enrolled and type of institution attended. A second set of questions addressed personal information, such as education of parents and siblings, whether they worked outside of school, among others. A third set of questions addressed participants’ perceived quality of education and confidence in future success in STEM disciplines, both academically and career-wise. Participants were asked to rate their education thus far on a 1-10 scale and indicate whether they felt insecure, neutral, or confident about performing well in STEM subject tests. A fourth set of questions included essays addressing their scientific curiosity and interest in science. Finally, we assessed the participants’ previous academic record in four STEM subjects: biology, chemistry, physics, and mathematics. High achievers were defined as those with scores equal to or above 80%, and low achievers as those with scores below 80%. Some participants had not taken a course on one or more of these subjects and were thus indicated as “no previous course”. Survey participants answered all questions via internet (www.clubesdecienciabolivia.com). The survey was not anonymous, however, the CdeCBo admissions committee only had access to basic demographic and essay questions (See Selection of Students). Participants were made aware of this and had the option not to respond to questions not evaluated by the admissions committee.

### Course design

Each course was designed by a scientist based in the United States or Europe in collaboration with a Bolivian investigator and both instructors taught the course together in Santa Cruz, Bolivia. Each course consisted of 10 hours of discussion-based lectures, and 20 hours of project-based laboratory work over the course of 5 days. After the instructional section of the course, students were required to present projects related to the course. Projects were decided by the instructors and included building homemade microscopes, creating a “cyborg” cockroach, performing genome engineering in bacteria, among others. (See Tables S1 and S2). The 6 Life Sciences courses analyzed in this study were: Model organisms and disease modeling (17 students); Microscopy and microscope design (17 students); Neuroscience and brain plasticity (21 students);Genome engineering and molecular biology (20 students); Microbiology and microbiome-host interactions (17 students); Regenerative Biology and diabetes (17 students). A description and a syllabus for each course are available in Table S1 and S2.

### Fundraising strategies

A major issue in developing countries is the lack of laboratory equipment at universities and research institutions [33]. Yet, several options exist in the United States to help scientists in the developing world, including universities’ equipment recycling programs, as well as non-profit organizations that administer equipment donations [33]. We took advantage of these programs and obtained several pieces of equipment, as well as laboratory supplies, from Harvard University.

While PhD students in the United States and Europe often have teaching assistantship requirements, they are rarely given the opportunity to design and execute their own course curricula. Importantly, PhD training grants in the United States include money to be used towards external training of students, including attendance to conferences and courses, which can be used to train students in teaching. We formed partnerships with Harvard University and Johns Hopkins University, which in return covered most of the costs associated with courses lead by their graduate students. These partnerships allowed us to implement 2 of the courses (Model Organisms and Neuroscience, respectively).

In Bolivia, on the other hand, our fundraising approach was different. Bolivian regulations require certain private companies and institutions to donate a percentage of their earnings to charity and nonprofit organizations. These donations can be made in cash or services (in-kind donations). As a registered foundation, we received several donations. While most donations were in-kind (flights, lodging, event planning services, or laboratory supplies), a small number of companies donated money that was used towards buying further equipment, as well as organizational costs of the program. Reagents and equipment were donated by the U.S. Embassy in Bolivia, Harvard University, Johns Hopkins University, and The Odin. Traveling costs of GAC and DGO were covered by their respective PhD programs. Additional foreign-based instructors’ flights were donated by Boliviana de Aviacion and the U.S. Embassy in Bolivia. Hotel rooms were in-kind donations. Laboratory and classroom space were provided free-of-charge by Universidad Privada de Santa Cruz de la Sierra. A complete list of all equipment and supplies, as well as their sources, is available in Table S3.

### Instructor selection and training

Foreign instructors were recruited by contacting PhD programs in the United States and Europe, as well as by word of mouth. Word of mouth included our network, our Facebook page “Clubes de Ciencia Bolivia”, and presentations at conferences. Foreign instructors were either PhD candidates or postdoctoral fellows. In the United States, graduate students become PhD candidates after passing their qualifying exam. The equivalent qualification in Europe is completion of a Masters program and enrollment in a PhD program. All foreign instructors were required to have had at least one teaching experience (for instance, teaching assistantships or other outreach programs) prior to teaching in Bolivia.

Local instructors were recruited by word of mouth. All local instructors had completed at least one year of Masters-level education. The majority of the local instructors were either PhD candidates or recent PhD graduates. All local instructors had at least one semester of teaching experience prior to teaching in our program. Foreign and local instructors were paired based on research experience and interest. Together, they developed the courses based on their area of expertise, complementing each other’s strength. Communication between instructors was done on their own time via video calls and emails. The organizers then approved courses after discussing the content in detail. Resources on teaching strategies were made available to all instructors.

Prior to the start of the course, we held a team retreat in Bolivia. This retreat was mandatory for all foreign and local instructors. In addition, we invited experts to further train and discuss with instructors on project-based learning, as well as challenges of teaching science in Bolivia.

### Student recruitment

We used three different channels to recruit students: press, social media, and visits to schools and universities. We opened applications on August 6th, 2017, Bolivian Independence Day, a major holiday and one of the top newspaper circulation days. We coordinated a simultaneous press release in the top three newspapers in the country: *El Deber* (Santa Cruz); *Los Tiempos* (Cochabamba) and *Página Siete* (La Paz). Each newspaper wrote, free of charge, a two-page article describing the program and inviting students to apply. Importantly, *El Deber* newspaper made this article a cover article. In addition, we coordinated appearances on several TV shows. Moreover, we orchestrated several other articles, including interviews to instructors, in different press media throughout the application period ending October 15th, 2017.

In social media, we created a Facebook page named “Clubes de Ciencia Bolivia”, which at the start of the application period had over 33,000 followers. Page viewing statistics show that our online presence reached over 45% of 18-24-year-old Bolivians with Internet access every week, suggesting that this approach was optimal to target a large audience. Working with local filmmakers, we created videos for recruitment, featuring instructors introducing their course, as well as multimedia presentations illustrating the potential for positive impact from scientific engagement in national development (see Video S1).

In order to recruit additional students, particularly in rural areas, we organized on-site visits to schools across the country. Our program had had two previous editions and at the time had over 300 alumni, several of which volunteer with the organization. These volunteers are based in 6 of 9 departments in Bolivia. Volunteers were trained online and given a standard presentation. They then visited schools and taught potential students how to apply.

In order to facilitate the recruitment of students, we incorporated an application platform online, which could be accessed via cell phones or computers. Cell phone penetrance in Bolivia is high. While no official numbers are available, a recent study in hospitals of the cities of La Paz and El Alto found that over 96% of patients ages 18-29 owned a cell phone [34]. Recent efforts from the Bolivian government have delivered cell phone service in rural areas of the country. Moreover, Quipus, a government-subsidized company has provided computers to school all around Bolivia. Altogether, we believe that the combination of press coverage, social media presence, and school visits provided an easily accessible application platform ensured the targeting a diverse student pool.

### Selection of Students

For this study, 113 students were selected from a pool of 903 eligible applicants. Only 109 students are part of the study, as 4 did not attend all sessions of their respective course. The gradable component of our online application form focused on essay questions rather than quantitative aspects (i.e. grades). Each application was reviewed and ranked by at least one member of the admissions committee. The admissions committee was blind to previous academic performance and emphasized representation of all corners of the country. Students were required to rank their top 3 courses of a total of 16 courses available to them. These included the 6 courses in biology-related fields analyzed for this study, as well as courses in computer science, engineering, entrepreneurship and social sciences. Students were considered part of the “Initial Interest” group if they were assigned to one of the 3 courses they selected. Students were part of the “No Initial Interest” group if they were assigned to a course different from their 3 selections. In total, 32 (29.36%) students were in the “No Initial Interest” group, while 77 (70.64%) students were in the “Initial Interest” group. Instructors were blind to which students were part of each group, although they were aware of the experiment. In order to allow for student preparation, the students were notified of their final course assignment at least 14 days prior to the start of the program.

### Student consent

The Clubes de Ciencia Bolivia Review Committee reviewed and approved the design and execution of this experiment and program. In accordance with Bolivian regulations, students were informed about the study prior to the application process. In order to participate in the program, students were required to consent to make pre- and post-test results available for anonymized publication. Upon admission, students had to submit two documents: a copy of an identification document proving their Bolivian citizenship and a letter from their high school or college proving that they were registered students and were in the year they indicated in the application. Occasionally, as this took place during school recess, we also accepted their last school report as proof. In the case of students under the age of 18, legal age in Bolivia, parents had to consent and send a signed and notarized authorization. Because a large number of students resided in different departments, a template letter was provided as part of the admission package.

### Testing

In each course, the instructors administered a 5-question test focused on the material to be covered in class lectures. Students were tested at the beginning and end of the course. Tests were graded by both instructors. Importantly, the pre-tests were not returned to the students before the post-test, nor did they have access to the answer key. Moreover, students were unaware that the pre- and post-tests were identical ahead of time. Each test was graded by each of the two instructors of the course. The average of both grades was then computed for analysis. A complete list of questions is available in Table S4.

### Statistics

For analyses that considered two groups of students, Two-tailed paired or unpaired t-tests were performed using PRISM version 7.0, depending on whether the groups being compared contained the same set of students (before vs. after the course) or different students (e.g. No Initial Interest vs. Initial Interest students). Gaussian distributions were assumed. Significance was set at a threshold of p < 0.05.

For analyses that considered three or more groups of students, we performed one-way ANOVA with Tukey’s correction using PRISM version 7.0. Gaussian distributions were assumed. Significance was set at a threshold of p < 0.05.

To compute the correlation between educational level, type of institution attended, previous academic achievement, or gender and confidence in success in science, we performed a multinomial logistic regression.

Finally, correlation analyses between two parameters were performed by computing Pearson correlation coefficients (Gaussian distributions were assumed) with PRISM 7.0.

## RESULTS

### Bolivian students rate their scientific education highly

To gain insight into the lack of relative representation of Bolivia in STEM fields, we conducted a country-wide online survey targeting precollege (11th and 12th grades) and college students in the context of an application to a science outreach program (see Methods). Respondents included students from 8 of 9 Departments in Bolivia, ages 15-22. A total of 903 students responded to the survey, which included questions designed to gauge participants’ perceptions of their educational experience, interest in science, and career goals. Notably, the sample group comprised 48.7% males and 51.3% females, equally spread across public (53.5%) and private (46.5%) institutions. Moreover, 30.0% of respondents were enrolled in high school, while 70.0% were in college (Table S5).

Surprisingly, respondents both at the precollege and college levels rated their educational experience highly (7.75 ± 0.08 precollege, n = 268 and 7.59 ± 0.05 college n = 614, on a 1-10 scale; p = 0.0736) (Figure 1A). Interestingly, while this observation was independent of gender (7.59 ± 0.06 n = 430 males and 7.69 ± 0.06 n = 451 females, on a 1-10 scale; p = 0.2433), respondents attending private institutions reported a slightly higher satisfaction level with the quality of their education (7.52 ± 0.06 public, n = 467 and 7.76 ± 0.06 private n = 406, on a 1-10 scale; p = 0.0067). Overall, our survey results indicate that Bolivians perceive their educational experience as positive, regardless of education level, gender, or type of institution attended (Figure 1A).

**Figure 1.**
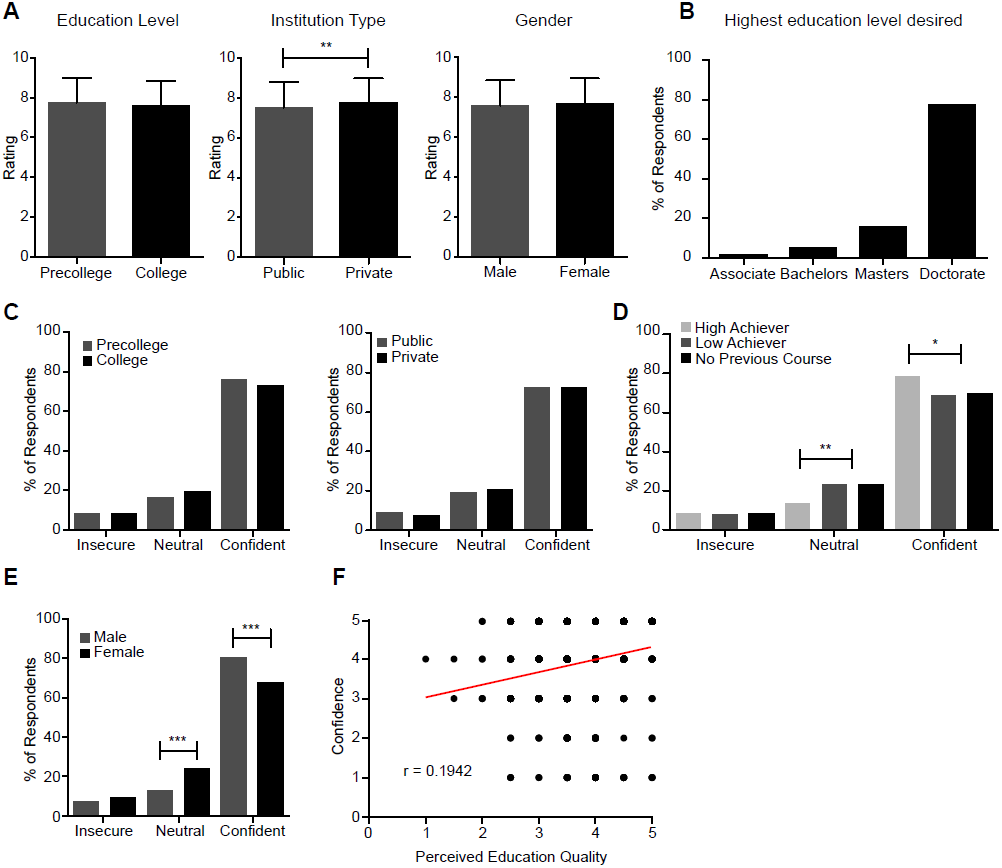
Perceptions, confidence, and aspirations of Bolivian youth in science. A) Perception of education quality does not depend on educational level or gender, but it is impacted by type of institution attended (**). Rating 1-10, with 10 being highest education quality. Bars represent mean ± SD. B) Highest educational level desired. C) Confidence in good performance in future STEM subject tests does not depend on educational level or type of institution attended. D) High achievers in biology tests (over 80%) are significantly less likely to be neutral (**) and more likely to be confident (*), compared with low achievers (under 80%) and those who had never taken a biology course. E) Confidence in good performance in future STEM subject tests is affected by gender: female respondents were more likely to be neutral (***), whereas male respondents were more likely to be confident (***). Neither gender was enriched in the group of respondents that felt insecure. F) Correlation between confidence in future good performance in STEM subject tests and perceived education quality (r = 0.1942). Each dot represents the response of a group of students with coordinates (perceived education quality, confidence). Unpaired t-test across groups in (A). Multinomial logistic regression when comparing more than two groups in (C-E). * = p < 0.05; ** = p < 0.01; *** = p < 0.001. N = 903

We then asked respondents to indicate what is the highest level of education that they would like to achieve, as well as whether they knew the requirements to pursue a career in science. Curiously, the vast majority (91.8%) of respondents reported to understand the required training to pursue a career in science (Figure S1A) and hoped to obtain a graduate level of education (77.6% Doctorate and 15.7% Masters, compared with only 5.2% Bachelors and 1.6% Associate level of education) (Figure 1B). These numbers are in great disagreement with current data showing that Bolivia has a vanishingly low number of researchers (<1 PhD-level scientist per 1,000 workers) [4, 10] or the fact that PhD-level faculty researchers in Bolivia are forced to undertake large teaching loads, diminishing their time for basic research [10].

In order to understand if our sample population was biased towards high academic achievers in the sciences, we requested respondents to voluntarily report their academic grade in their previous biology courses. Of the 876 respondents who opted to report their grades, 23.3% had never taken a biology course, whereas 25.6% had but achieved a low grade (<80%) and 51.1% obtained a high grade (>80%) (Figure S1B). Similar numbers were obtained when inquiring about courses in Chemistry, Physics, and Mathematics (Figure S1B). Of note, no strong correlation (r < 0.5) between academic achievement in any of these subjects and perceived education quality was found (Figure S1C).

### Bolivian students have confidence in their scientific aptitude

To gain further insight into students’ perceptions and attitudes towards STEM careers, we asked respondents how confident they were in achieving a high score in a STEM subject test. We found the majority of respondents to be confident that they will perform well in STEM courses (Figures 1C-E). Notably, no clear distinctions were observed in confidence levels when participants were grouped by education level or type of institution attended (Figure 1C). Nevertheless, confident participants were more likely to be high academic achievers than low academic achievers or have not taken a biology course previously, whereas neutral respondents were the least likely to be high achievers (Figure 1C). Interestingly, male participants reported higher levels of confidence than their female counterparts (Figure 1D). Finally, we found no clear correlation (r = 0.1942) between confidence in future STEM subject tests and perception of education quality (Figure 1E). Altogether, our results show that the Bolivian student population perceives their education as being of high quality, intend to pursue advanced science degrees, and are confident that they will succeed in STEM subjects.

### Life Sciences workshops effectively transmitted new concepts

Given the surprising results obtained in this initial nation-wide survey, we decided to test the knowledge of a subset of respondents who we admitted in our science outreach program, “Clubes de Ciencia Bolivia” (CdeCBo), in 6 life sciences topics, namely model organisms, microscopy, neuroscience, genome engineering, microbiology, and regenerative biology. During the selection process for our program (described in Methods) the committee in charge ensured that the program represented the country’s demographics (Figure 2A). Overall, 8 of 9 departments in Bolivia were present in the student body. Due to possible migration within Bolivia, we classified the students based on their department of residence, not department of birth. Nevertheless, only 16 of 109 (14.68%) students were residing in a department different from their department of birth (data not shown). Of note, one department, Pando, was not present in the application pool and therefore could not be included in the study. In addition, 3 departments, Potosi, Tarija and Chuquisaca, were underrepresented in the applicant pool and thus in the student cohort (Figure 2A). The admitted student pool displayed an equal representation of gender, type of institution in which the students were enrolled, and a balanced distribution of age (Figure 2B-D). Because the application was open to students from 11th grade in high school to third year of college at the time of application, the student cohort was enriched for college students over high school students (Figure 2E). Importantly, because the experiment was performed during summer vacation, some students had graduated high school but had not yet started college. Together with the high school students, we categorized this group as “precollege”. Moreover, the committee gave preference to applicants who had not attended our program in past years (77 of 109 participants), diminishing the possibility of any advantage gained from previous exposure to our approach that might confound our data (Figure 2F). In fact, as 2 of 6 courses had been taught in previous years, the returning students were not allowed to attend the same course twice. Altogether, to the best of our possibilities, our study analyzed a cohort of students well balanced with regards to age, gender, previous education quality and level, as well as representative of the demographics of Bolivia. Each course was taught by 2 instructors: 1 instructor based in a foreign country (United States or Europe) and 1 local instructor.

**Figure 2.**
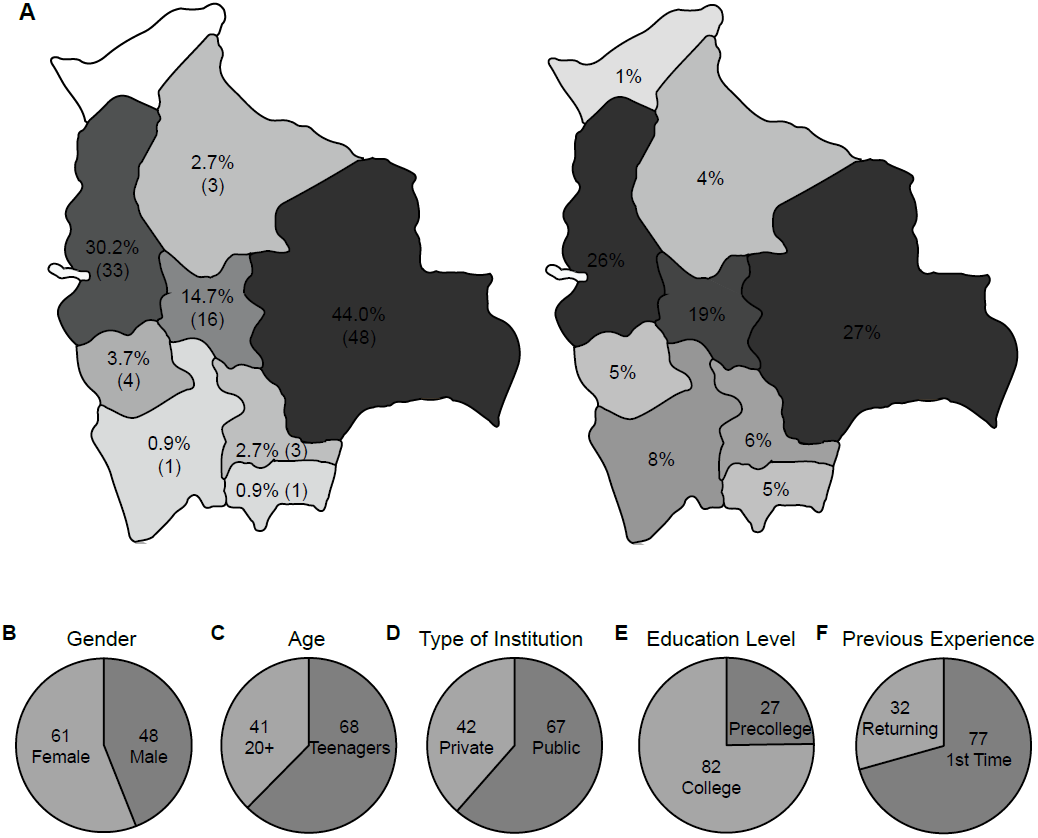
Demographic distribution of the student cohort. A) The student body represents the population of Bolivia. Left: Map of Bolivia showing the number of students from each department represented in the student cohort, as well as the corresponding student distribution (%). Right: Population distribution (in %) in Bolivia according to the most recent census (2012). For easy visualization, maps were colored in shades of gray reflecting the relative population distribution. B-F. Other demographic parameters studied: B) Gender, C) Age, D) Type of institution in which the students were enrolled, E) Education level of the students, F) Previous participation in non-competitive science outreach programs. N = 109.

The courses contained a theoretical component, taught using a discussion-based lecture approach (Figure S2A), as well as a project-based laboratory component (Figure S2B) [35, 36]. At the end of the courses, all students were required to present on a topic within the scope of their course that they researched in small groups of 3-4 students during the week (Figure S3). We designed questions to assess comprehension of basic principles behind biological processes, such as heredity, immune rejection, or neural communication (Table S4). As we aimed to understand whether students had a basic grasp of these topics, we designed questions targeting knowledge and comprehension, following the principles detailed in the Bloom’s taxonomy of questions [37]. Strikingly, we found that incoming students performed poorly in our tests (38.24%±20.42, N = 109) (Figure 3A), revealing a stark disconnect between perceived education quality and aptitude in STEM subjects.

**Figure 3.**
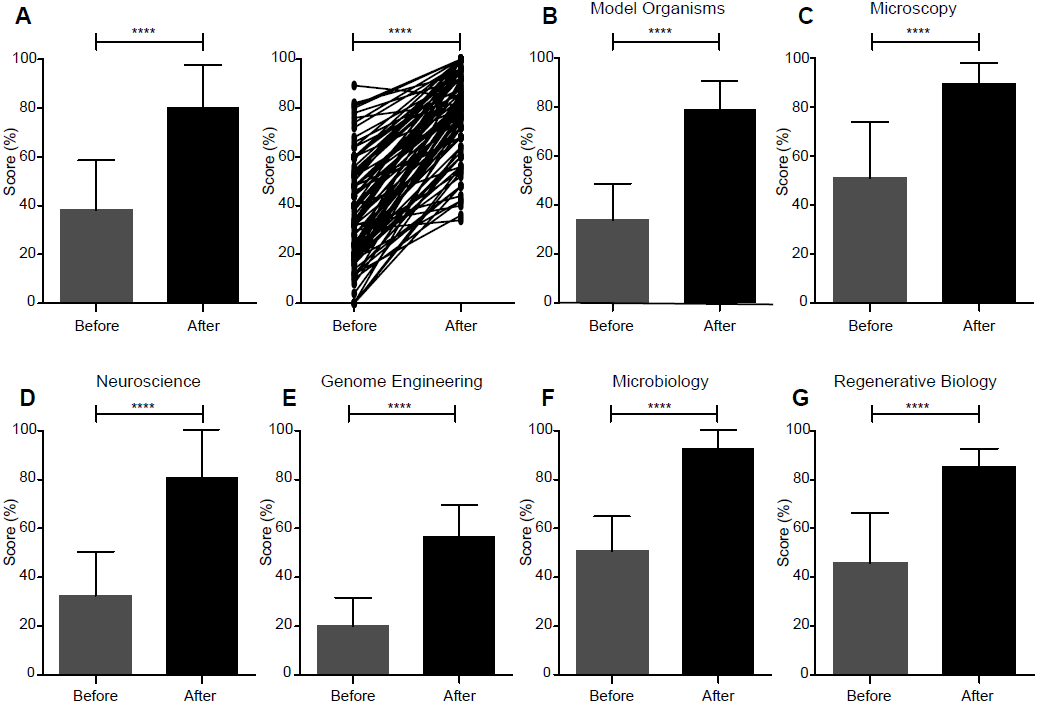
Effective transmission of key concepts in biology. Comprehension of specific topics was evaluated before and after the courses. A) Overall student performance before and after all 6 courses. Left: Combined student performance. Right: Paired test results for each student. N = 109. Paired t-test; **** = p < 0.0001. B-G) Student performance for each individual course. B) Model organisms and disease modeling. N = 17. C) Microscopy. N = 17. D) Neuroscience. N = 21. E) Genome Engineering. N = 20. F) Microbiology. N = 17. G) Regenerative Biology. N = 17. Bars represent mean ± SD. Paired t-test; **** = p < 0.0001.

In order to interrogate the effectiveness of our approach to teaching biology, we asked whether taking a course increased the score in the same tests. Overall, test scores after taking the courses were significantly higher (N = 109, p < 0.0001) than those obtained at the beginning of the program (Figure 3A). In fact, 108 of 109 students increased their test score after participating in the program (Figure 3A). Of note, this pattern was true for every individual course analyzed (Figure 3B-G), from the course with the lowest test scores, Genome Engineering (Figure 3E, n = 20 p < 0.0001) to the one with the highest scores, Microbiology (Figure 3F, n = 17, p < 0.0001), demonstrating that the efficiency of our approach extends beyond a particular topic or instructor.

### Possible confounding factors did not influence students’ performance

Next, we investigated whether our results were confounded by external factors. Importantly, we found no influence of gender or age on test scores obtained either before or after taking a course (Figure 4A-B). Curiously, however, students coming from public institutions scored slightly better than those enrolled in private institutions before taking the course (p < 0.05), yet no difference was found in scores after taking the course (Figure 4C). In Bolivia, public universities tend to be higher ranked than their private counterparts [38]. Therefore, the statistically significant difference in scores between students enrolled in public versus private institutions is likely to have been driven by college students, who outnumbered precollege students in our program (Figure 2E).

**Figure 4.**
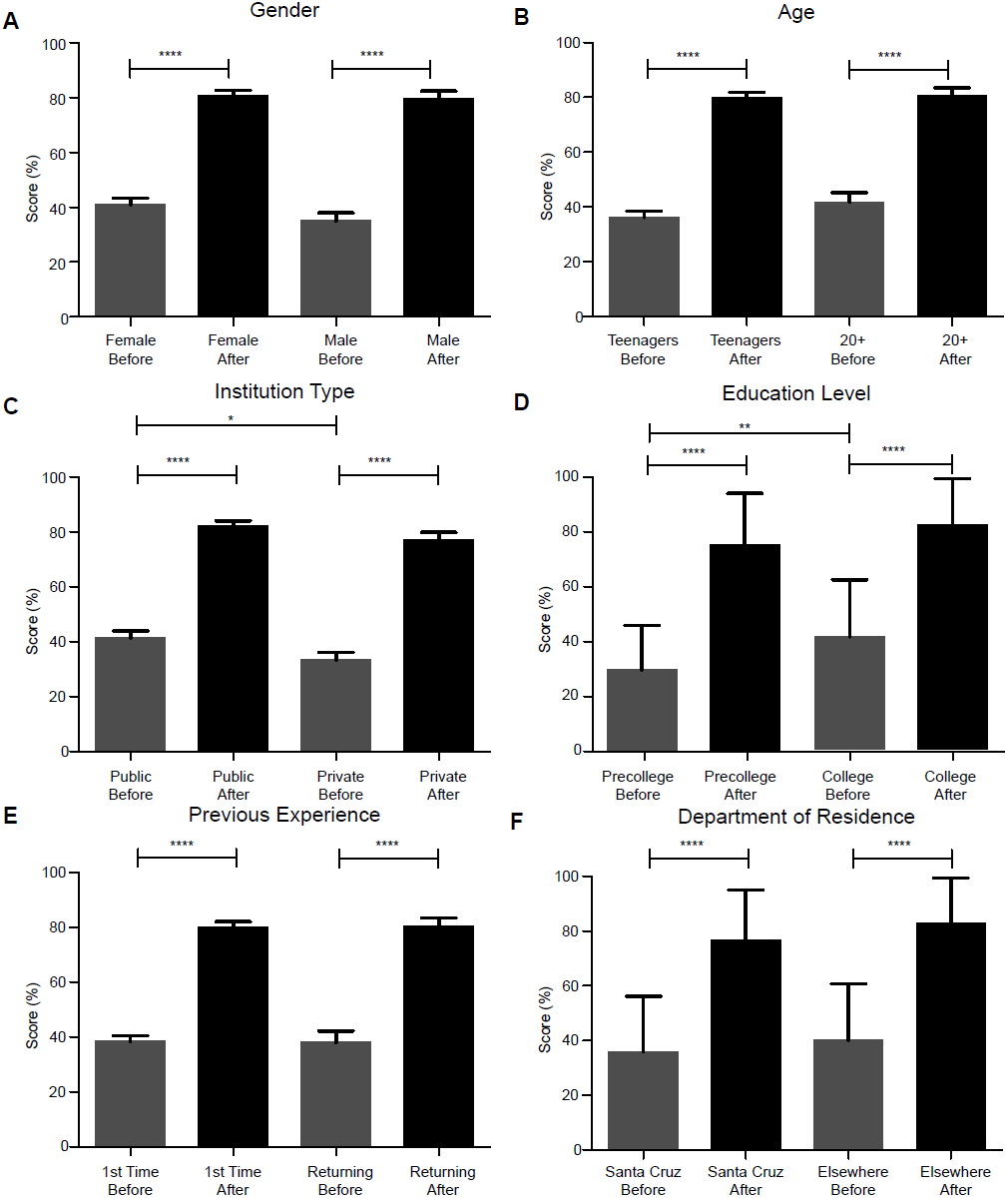
Confounding factors do not affect student performance. Comprehension of specific topics was evaluated before and after each course. Test scores after each course were not significantly affected by any of the following possible confounding factors. A) Gender. Female n = 61, Male n = 48. B) Age. Teenagers n = 68, 20+ years old n = 41. C) Type of institution in which students were enrolled. Private n = 42, Public n = 67 (unpaired t-test Public test score before course vs. Private test score before course, *). D) Education level of the students. Precollege n = 27, College n = 82. E) Previous experience in non-competitive outreach programs. First time in an outreach program n = 77, Returning student n = 32. F) Department of residence. Santa Cruz n = 48. Elsewhere n = 61. Bars represent mean ± SD. N = 109. Paired t-test within group, unpaired t-test across groups; * = p < 0.05; ** = p < 0.01; **** = p < 0.0001.

As expected, college students performed better than precollege students in the test before taking the course (p < 0.01). Still, no difference was found in test scores after taking a course, indicating that education level did not impact the extent of learning (Figure 4D). This observation is in agreement with previously published data showing that students from different backgrounds can achieve the same learning gains in project-based biology courses [36]. Moreover, these data are also consistent with the notion that project-based STEM courses “level the field” by preferentially benefiting students at an initial disadvantage [35].

One could argue that previous exposure to similar outreach programs could confer an advantage in taking the tests used in this study. As there are no other major non-competitive science outreach programs in Bolivia, as opposed to science olympiads, for example, we only accounted for participations in our initiative in previous years, 2016 and 2017. Comparison of test scores between first time students and those with previous experience revealed no significant difference in test scores either before or after the course between these two groups (Figure 4E), suggesting that our approach can be used to educate student populations independently of their previous access to similar opportunities. Of note, as mentioned in Methods, returning students were not allowed to retake the same course from previous years.

Bolivia is a very diverse nation, with large disparities in income and access to education between different departments [24, 39]. Abundant literature shows strong correlations between socioeconomic status and educational attainment [40]. Santa Cruz, in particular, is the wealthiest department in Bolivia *per capita*, and students residing in Santa Cruz are the most represented group in our study (48/109). Nevertheless, our results demonstrate that there is no significant difference in scores either before or after the courses between students residing in Santa Cruz and those coming from other departments (Figure 4F). However, we did find a modest but statistically significant (p < 0.05, N = 96) decrease in the post-course test scores of students who reported to work after school in comparison with those who did not (Figure S4A). Work obligations from a young age, prevalent in rural parts of Bolivia, have been shown to result in slower learning speeds, disinterest in academics, and ultimately higher school dropout rates [24]. It is thus possible that this small decrease observed in our study hints at differences in learning ability not captured when not specifically sorting out students who work while pursuing their studies. Nonetheless, none of several other possible confounding circumstances impacted test scores after the courses. In addition to those mentioned above, we also found no significant differences after dividing students based on the highest academic degree they desire to acquire (Figure S4B), or their English skills (Figure S4C), education level of their parents and siblings (Figure S4D), perceived education quality (Figure S4E), previous academic success in biology classes (Figure S4F), confidence in performance in STEM subject tests (Figure S4G), previous exposure to STEM professionals (Figure S4H), or even preferred mode of acquiring new information (visual or verbal; Figure S4I).

### Initial interest in the course material does not affect learning

Next, we sought to determine whether the extent of learning depended on interest on the specific topic taught in a course. Interest in a topic is well documented to influence motivation and scholar achievement related to that topic [41]. Conversely, becoming more skilled and knowledgeable about a topic tends to increase an individual’s interest on it due to increased positive associations [42]. To investigate the possibility that previous interest in a course altered the extent of learning after participating in that course, we compared test scores between students who were placed in courses outside their list of preferred options (No Initial Interest) with those who attended a course in their preferred field (Initial Interest). Strikingly, students in the “No Initial Interest” group did not perform differently from those enrolled in a course on their list of three top choices (i.e. “Initial Interest” group) across all 6 courses (Figure 5A-F). Students from either group attained equally significantly higher test scores after taking their respective courses (p < 0.0001), suggesting that our model of science teaching was effective even to those students who were not directly interested in the specific subject taught *a priori*.

**Figure 5.**
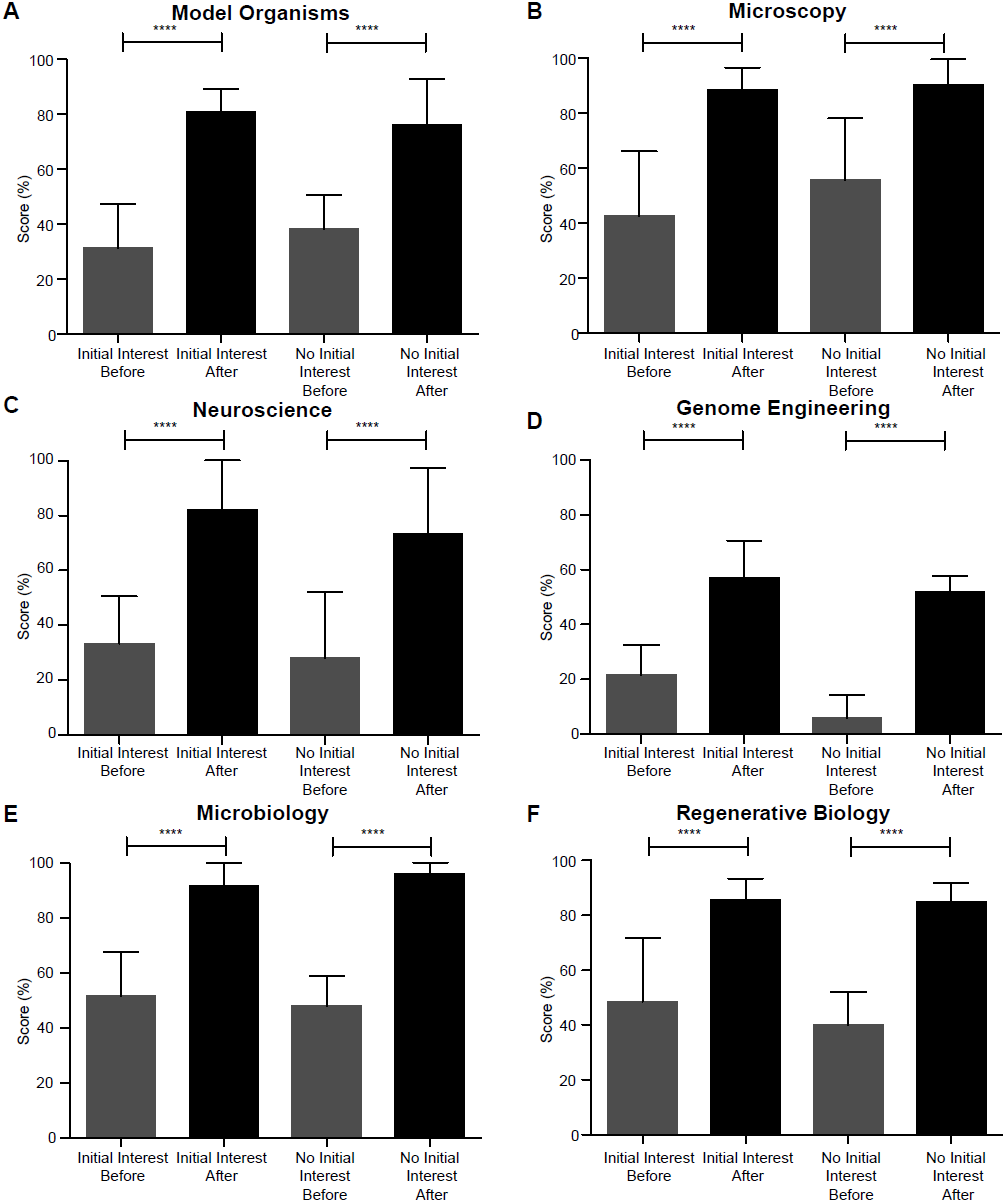
Initial interest in the topic does not affect student performance. Comprehension of specific topics was evaluated before and after each course and the data was divided into two groups according to students’ interest: Initial interest and No initial interest. Test scores were not significantly different between these two groups. A) Model organisms and disease modeling. Initial interest n = 11; No initial interest n = 6. B) Microscopy. Initial interest n = 6; No initial interest n = 11. C) Neuroscience. Initial interest n = 18; No initial interest n = 3. D) Genome Engineering. Initial interest n = 18; No initial interest n = 2. E) Microbiology. Initial interest n = 12; No initial interest n = 5. F) Regenerative Biology. Initial interest n = 12; No initial interest n = 5. Bars represent mean ± SD. N = 17-21. Paired t-test; **** = p < 0.0001.

In summary, the lack of significant differences in the final, i.e. after taking the course, test scores between genders, age groups, previous exposure to science outreach, education level of the students or their families, previous academic success in biology classes, department of residence, or initial interest in the specific course topic, amongst others, strongly suggest that our approach to teaching biology in a dynamic and inquisitive way may be broadly applicable in other developing countries.

### Hands-on workshops positively impact students’ enthusiasm for STEM

Finally, we sought to assess the students’ satisfaction with our program and characterize their own personal attitudes towards science. When asked to rate different aspects of the CdeCBo initiative upon its completion, the vast majority rated their experience very highly (4 or 5, in a 1-5 scale, with 5 being most satisfied), with regards to having both a foreign and a local instructor teaching together (Figure 6A), course activities (Figure 6B) and materials (Figure 6C), motivation of the instructors (Figure 6D), clarity of course objectives (Figure 6E), and connection between theory and practice (Figure 6F). These are important data, as the teaching methods utilized by us are vastly different from those used in Bolivia for science education, which could have caused attrition and dissatisfaction amongst students. Furthermore, the students also reported high levels of excitement for science at the end of the program. When asked how strongly they agreed that science is exciting, enjoyable to participate in, a possible career choice, or whether they enjoy solving scientific problems and that hard work will help them succeed in science, most students responded 5 in a 1-5 scale, with 5 being strongest agreement (Figure 7A-E). In additional questions, most students stated they would like to participate in similar programs in the future (Figure 7F) want to keep learning more about science (Figure 7H), definitely want to become scientists (Figure 7I), and thought they could succeed in science (Figure 7J). More importantly, 89 of 91 students stated that our program increased their interest in science (Figure 7F).

**Figure 6.**
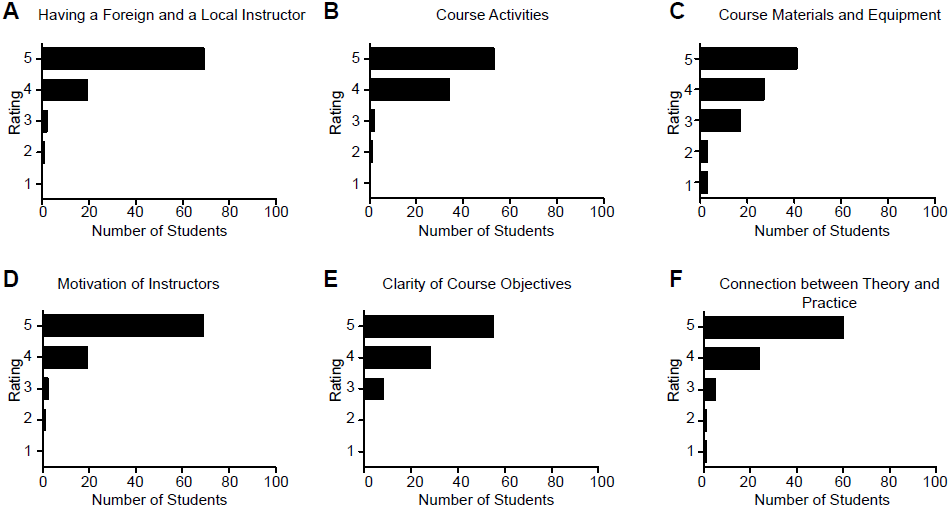
Students report a very positive experience after participating in our initiative. Students were asked to rate several aspects of our initiative. A) Having a foreign or a local instructor. N = 109. B) Course activities. N = 90. C) Course materials and equipment. N = 91. D) Motivation of instructors. N = 91. E) Clarity of course objectives. N = 91. F) Connection between theory and practice. N = 91. Rating 1-5, with 5 being most satisfied.

**Figure 7.**
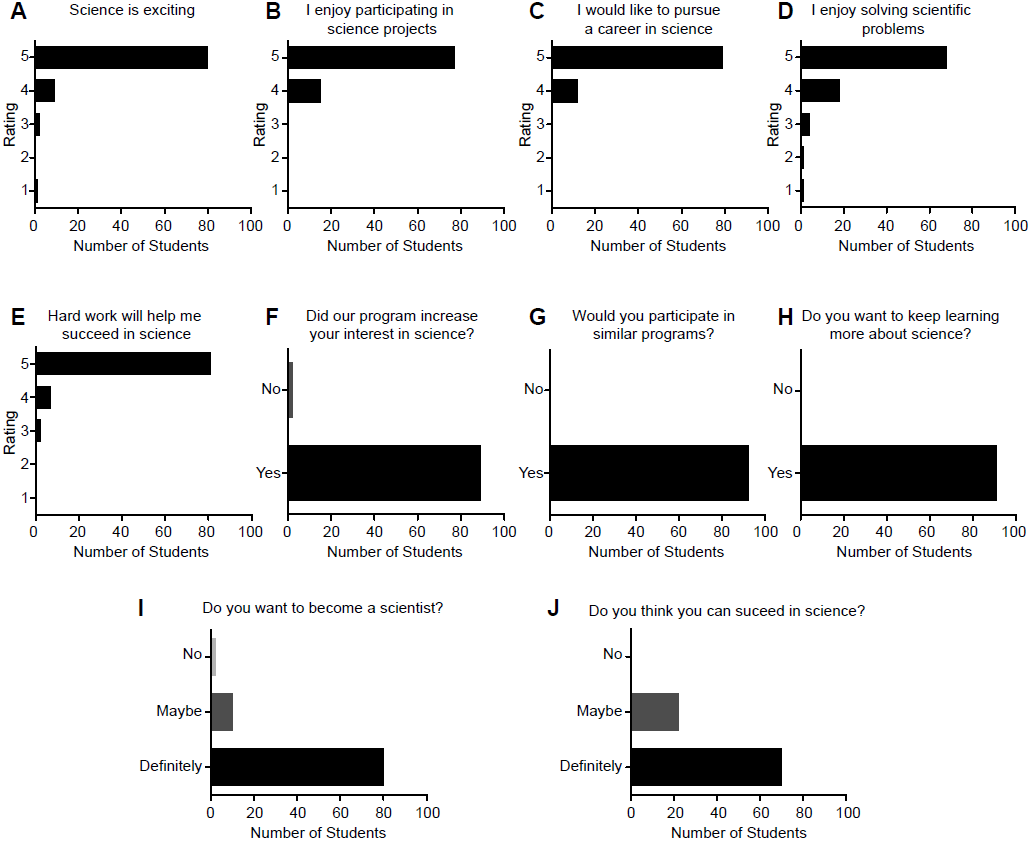
Students report positive attitude and increased interest in science after participating in our initiative. Students were asked several questions pertaining to their interest in science. A-E) Students gave a rating 1-5, with 5 being the strongest agreement with the question posed or statement made. A) Science is exciting. N = 92. B) I enjoy participating in science projects. N = 92. C) I would like to pursue a career in science. N = 91. D) I enjoy solving scientific problems. N = 92. E) Hard work will help me succeed in science. N = 90. F-H) Students replied Yes or No to the posed question. F) Did our program increase your interest in science? N = 91. G) Would you like to participate in similar programs? N = 92. H) Do you want to keep learning more about science? N = 92. I-J) Students replied No, Maybe, or Definitely to the posed question. I) Do you want to become a scientist? N = 92. J) Do you think you can succeed in science? N = 92.

To sum up, students in Bolivia demonstrated thorough satisfaction with their education, high confidence in their scientific skills, and a widespread desire to pursue advanced scientific degrees. Even though initial low test scores before taking our courses may have challenged that self-perception, students acquired new knowledge in a teaching methodology new to them, intensive hands-on biology courses, in a manner independent from most possible confounding factors, and reported increased interest in science after completing the courses.

## Discussion

Developing countries, and Bolivia in particular, have long been in the backwaters of research and technological innovation, despite a sizable population and GDP [11, 38]. It is not trivial to devise and implement measures that can position developing countries on par with developed nations with a history of prolific research and development [19, 20]. Notwithstanding, we envision that our efforts to transform the landscape of science education in Bolivia may help reverse this trend. Using biology courses as examples, we demonstrate that project-based instruction can be successfully implemented in a developing country and can adequately relay important biological concepts to local high school and college students of diverse backgrounds in a limited time span.

Studies in project-based learning have shown that the critical thinking of students exposed to this methodology is significantly increased over time [43]. Yet, not all students transition equally towards project-based learning, and students lacking experience with this approach can experience difficulty in its implementation [44]. Previous studies have shown that this teaching method strongly favors academically “low achievers” [45]. “High achievers”, on the other hand, tend to have a difficult transition towards project-based learning, with many of them abandoning the projects altogether [45]. Our student cohort was an interesting group; anecdotally, we observed that our application process recruited academically high achieving students from across Bolivia. The majority of our students have had little, if any, previous experience in project-based learning, as this methodology is not part of the Bolivian curricula. Yet, none of our students abandoned their course and 108 of 109 did better in the post-tests (Figure 3). These results not only challenged our expectations, but also support the importance of performing “on the ground” work across the globe. To date, most project-based education studies in underrepresented populations have left behind the developing world [28, 29, 46]. Yet, even comparisons of US-born Latinos to first generation Latino immigrants in the United States have consistently shown differences in several behaviors [47, 48], suggesting that underrepresented minorities in the developed world are not necessarily representative of their native populations, and highlighting the need of further studies in the developing world itself. In this study, we chose Bolivia for its high population diversity and striking economic disparity, but we foresee the need of performing similar projects in other regions of Latin America and across the developing world.

Investment in education at the early stages produces long-term benefits in decision-making [49]. Yet, this investment needs to be accompanied with policies that focus on training teachers on innovative approaches to education. Beyond financial limitations, developing countries have cultural and infrastructural issues that need to be solved in order to promote a better environment for education. For example, an observational study in rural Tarija, the most Southern region of Bolivia, registered several concerning behaviors in teachers that would be considered unacceptable in most other countries: favoritism, lack of interest, and unexcused absence from the classroom [24].

Poor academic performance and low student retention in Bolivia could be also attributed to child labor. In Bolivia, children have a strong sense of responsibility towards their family, and as such they enter the labor market at an early age [24]. Previous studies have shown that, in Bolivia, child labor reduces educational attainment by two years on average [50]. In 2014, the Bolivian government, disregarding international labor conventions, passed a controversial law reducing the legal working age to 10 years of age if self-employed and 12 years of age for contract work [51]. Recently, it has been shown that school dropout rate is positively correlated with a decrease in the minimal legal working age [52]. In line with this observation, Bolivia is the only country in Latin America where the percentage of students that drop out from higher education institutions is greater than those of still enrolled students and of those who completed their degrees, according to data collected by the World Bank [18].

We believe that addressing these intricate issues will require not only domestic changes in Bolivia, including policy changes and building new infrastructure for STI, but also collaboration with scientists abroad working in more established and vibrant research environments. Our initiative is unique in that it allows PhD students and postdoctoral fellows carrying out cutting edge research in the United States and Europe to collaborate with local Bolivian scientists primarily fulfilling teaching roles to design and develop project-based courses. Moreover, it allows considerable numbers of students from developing countries to meet and interact with scientists in a supportive, informal, and discovery-oriented environment over the course of five days, fostering the transmission of knowledge and experiences and, ultimately, the creation and nurturing of a local pool of talent. Furthermore, maintaining the course sizes small (17-21 students) leads, in our experience, to a close and lasting mentorship relationship between instructors and students, which extends well beyond the timeline of the courses. Together, these characteristics not only powered the efficacious transmission of scientific knowledge, but will also aid in developing and strengthening science in Bolivia in the medium and long term [29].

In our opinion, the research universities sponsoring this type of initiatives also benefit from their involvement in tangible ways. Having a graduate student teaching a course abroad allows universities to target two unique pools in recruitment: students in the developed world who are interested in teaching, and students in the developing world who are interested in becoming graduate students in the United States or Europe.

The current study has some limitations. First, it is not possible to completely account for selection bias in an initiative with eligibility and selection criteria that enrich our sample for students with high academic achievement and an interest in STEM. Nevertheless, we addressed this concern by categorizing students based on their science education perspectives, previous academic achievement, and initial interest in their course, amongst others, and saw no difference in post-course test scores (Figure 4, 5, S4). In addition, comparing students of two different education levels, precollege versus college, revealed no difference in post-course test scores (Figure 4D). In addition, because the format of the courses included lecture and laboratory components (Table S2), it is difficult to disentangle the direct contribution of each component to overall learning. Future studies including lecture-only and laboratory-only courses will be needed to further understand the impact of each component on learning.

Importantly, an unexpected limitation we encountered was more social in origin. Following CdeCBo 2017, we were surprised to see particular segments of national media and university professors cautioning against the adoption of modern science education techniques and the intellectual challenges these impose, despite overwhelming positive feedback from student participants and most independent media outlets. In 2017, one publication lamented CdeCBo’s forward-thinking approach in relation to the Bolivian “national patriotic agenda”:

> “The issues raised may be challenging and the future is always a challenge, but one should not lose sight of the present. In a country like Bolivia, where we have not even solved the basic health problems, we do clubs that require one to learn about the post-genome stages. [In] a stretch in which we are not manufacturers of bicycles, we think about principles of flight and aeronautical design…” [53].

The article goes on to note CdeCBo’s lack of emphasis on “ancestral knowledge,” and ponders whether CdeCBo’s neuroscience course, presenting modern experimentation techniques, should be “*reoriented toward primary education, infant health, to students’ cognitive processes, or to potentiating these to contribute to development of the country.*” The following year, following CdeCBo 2018, one print newspaper stressed the need to consider “national reality” in contrast to CdeCBo’s vision to “*practice a science that is academic, technocratic, prize-winner of modernity, not originated in the* [*2010 Bolivian education law*] *070.*”

While we fundamentally disagree with the assertion that exposing students to knowledge of modern scientific advancements will negatively impact Bolivia’s future, the point is well taken of a need for regional sensitivity and consideration for local contexts. An illustration of CdeCBo’s regional considerations comes in topics chosen during course design. Strikingly, some of the above authors performed a study of specific educational desires among Bolivian students in two distinct Bolivian departments, Cochabamba and Chuquisaca, reporting that the most highly sought topic pertained to HIV/AIDS education [54]. Indeed, one of CdeCBo’s most popular courses deals with genome editing in the context of immunological resistance to HIV (Table S2). Moreover, a publication from the same group went on to posit that “*science clubs* [*Bolivia*] *could encourage that the positive image about science and its consequent practice are not contradictory, but continuous… This activity will create a good basis for medical practice to combine assistance with solution, research with intervention.*” [55]. It is our view that many negative perceptions, among limited segments of population, toward modern scientific education approaches by CdeCBo can be alleviated through enhanced transparency and dialogue between these groups and CdeCBo instructors. We are confident that both camps have in mind Bolivia’s best interests for the future, and that open channels of mutually respectful dialogue will identify the most effective and beneficial means of national development.

To the best of our knowledge, no initiative similar to ours exists yet in the region. Future directions thus include carrying out longitudinal studies to evaluate our students’ future success in academic applications and their career choices in STEM versus other fields. In particular, it will be fascinating to investigate whether students who were allocated to courses that did not initially interest them will be any less likely to follow STEM careers than those who attended their preferred courses. Even though test scores were very similar (no statistically significant difference) between these two groups, only a long term follow-up study can shed light on whether their experience overall was similarly inspiring. In either scenario, exposure to biological research during the program has been shown to increase the participants’ odds of successfully applying to internships and scholarships to study biology at the college or PhD level [56, 57]. Ultimately, we aim to provide an experience which empowers students with the self-confidence, excitement, and participatory motivation to help them realize that they too can pursue successful careers as scientists and thrive alongside scientists of any other nation.

## Disclosure Statement

The authors reported no potential conflict of interest.

## Acknowledgments

The authors would like to thank all volunteers and staff members of the Clubes de Ciencia Bolivia Foundation, as well as the Universidad Privada de Santa Cruz de la Sierra (UPSA) for all their support.

## Author Contributions

LIRB, OGC and MAMR designed the study. LMRF, GAC, YR, DGO, NMR, DKAG, AV, SB, LGA, CS, and CVAS taught the courses and collected the data. LMRF, BLV, GAC, GVD, LIRB, and MAMR analyzed the data. LMRF, GAC, and MAMR wrote the manuscript with contributions from all authors.

---

**Figure S1. Survey respondents’ perceptions and academic achievement in science.** A) Majority of respondents reported to understand the requirements of pursuing a career in science. B) Academic achievement in previous science courses: biology, chemistry, physics, and mathematics. C) Success in previous science subject tests – biology (r = 0.2524), chemistry (r = 0.2703), physics (r = 0.3148), and mathematics (r = 0.3180) – weakly correlated with perceived education quality. Each dot represents the response of a group of students with coordinates (perceived education quality, score in previous science course). N = 903

**Figure S2. Images illustrative of the lecture and laboratory components of the courses.** A) Representative images of the lecture component of the courses. B) Representative images of activities performed by the students in the laboratory component of the courses.

**Figure S3. Images illustrative of the presentations by the students at the end of the courses.** Representative images of students preparing and presenting their work at the science fair held at the end of the courses.

**Figure S4. Additional potential confounding factors did not visibly affect test scores.** Comprehension of specific topics was evaluated before and after each course. Test scores were compared between groups of students divided by the following characteristics. A) Afterschool work. Doesn’t work n = 88; Works n = 8 (unpaired t-test Doesn’t work test score after course vs. Works test score after course, *). N = 96. B) Degree wanted. Bachelors n = 5; Masters n = 8; Doctorate n = 82. N = 95. C) English skill level. Basic n = 15; Intermediate n = 46; Advanced n = 34. N = 95. D) Family education. No siblings with a college degree n = 41; At least one sibling with a college degree n = 55; No parent with a college degree n = 19; At least one parent with a college degree n = 77. N = 96. E) Perceived education quality. Satisfied (rating ³ 8) n = 53; Not satisfied (rating < 8) n = 43. N = 96. F) Academic success in biology. High achiever (³80%) n = 65; Low achiever (<80%) n = 11; No previous course n = 20. N = 96. G) Confidence in science performance. Insecure n = 6; Neutral n = 9; Confident n = 81. N = 96. H) Exposure to STEM professionals. Little exposure n = 8; Some exposure n = 47. A lot of exposure n = 41. N = 96. I) Type of learner. Visual learner n = 68; Verbal learner n = 28. N = 96. Bars represent mean ± SD. Paired t-test within group, unpaired t-test across groups; * = p < 0.05; ** = p < 0.01; *** = p < 0.001. C.D., college degree. STEM, Science, Technology, Engineering, and Mathematics.

**Table S1. Course summaries**

**Table S2. Course syllabi**

**Table S3. Course materials**

**Table S4. Questions tested in each course**

**Table S5. Nation-wide survey demographics**

**Video S1. Recruitment video for Clubes de Ciencia Bolivia (CdeCBo) 2018.** The video begins with an artistic rendition of progress in science and the importance of exploring and connecting new ideas, followed by short clips of instructors, course activities, and the city where CdeCBo 2018 would take place, ending with information on how to apply. Created by local filmmakers in collaboration with members of our initiative. English subtitles have been added for this manuscript.

