## Supplementary figures and images for "Perceptions and prospects in life sciences in a heterogeneous Latin American population"

### Figure S1

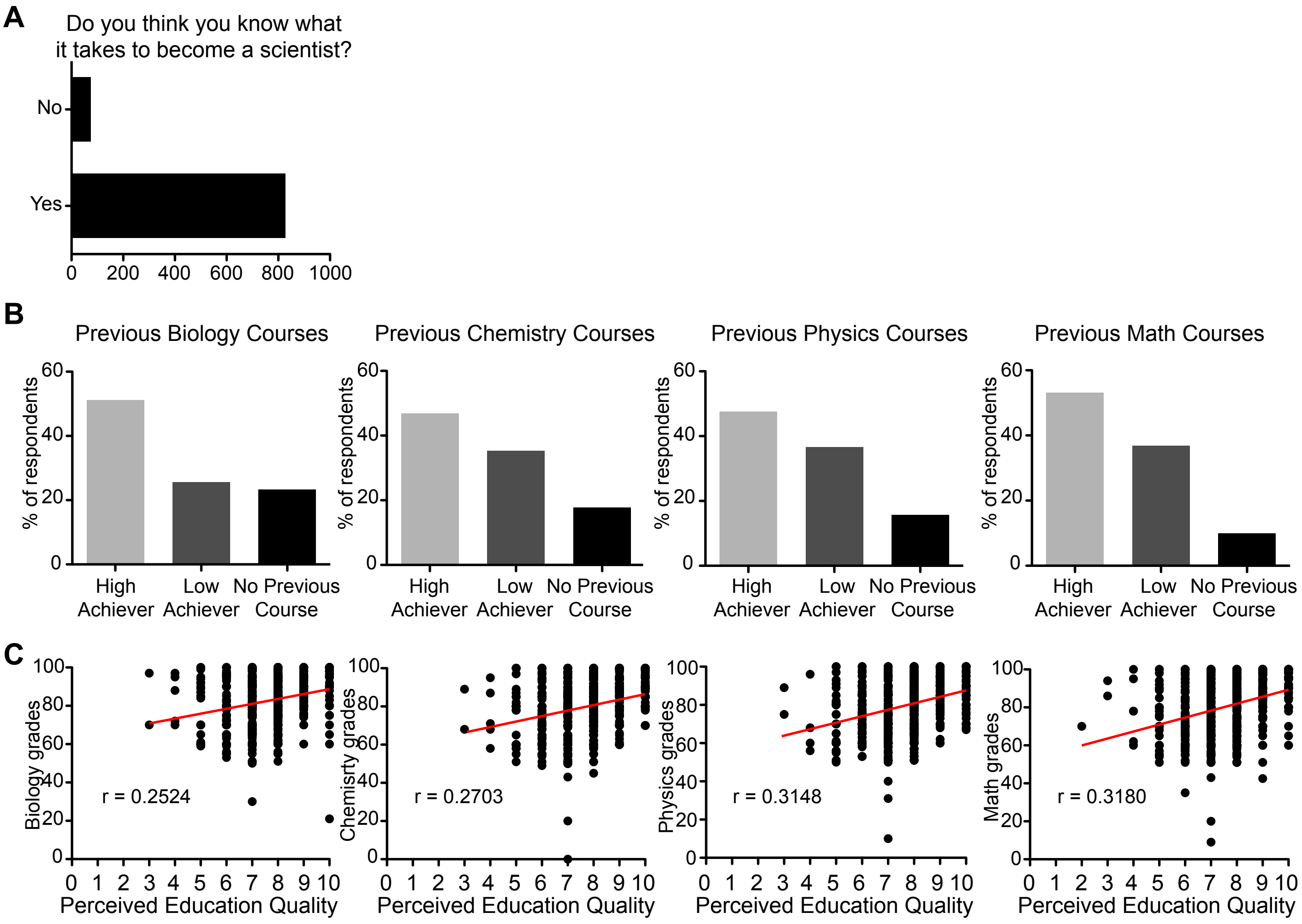


Ferreira et al., Figure S1

### Figure S2

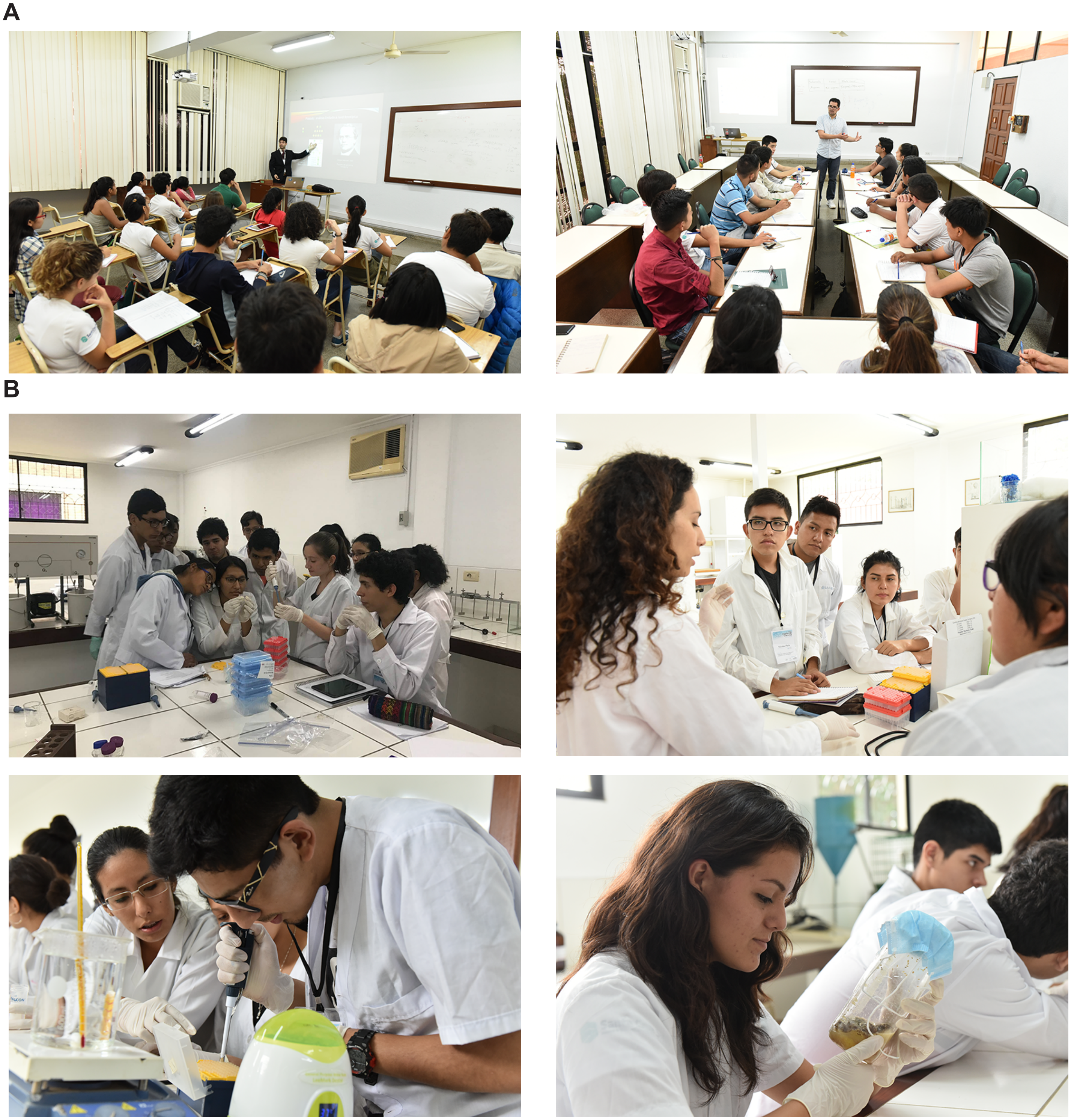


Ferreira et al., Figure S2

### Figure S3

A

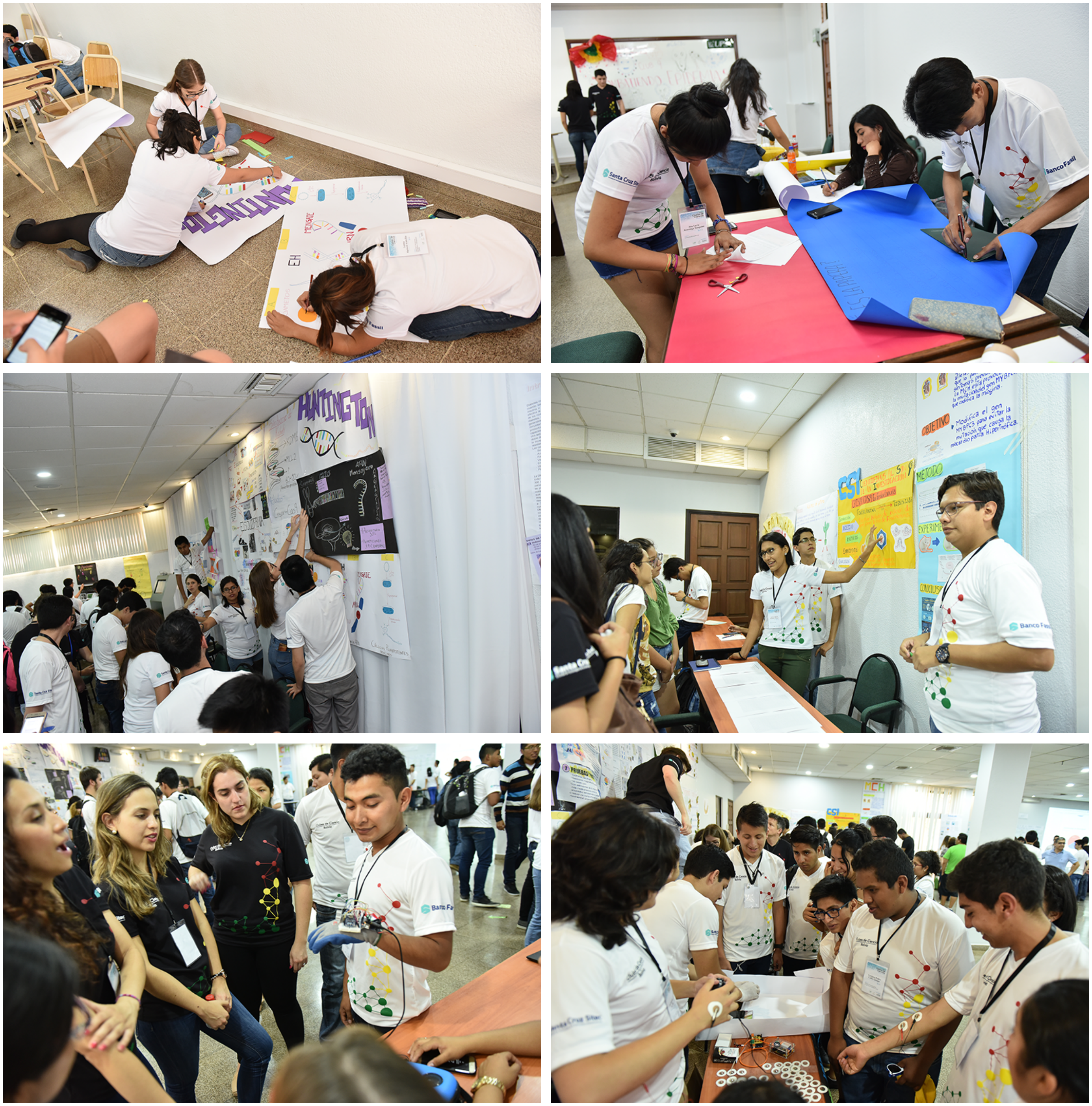


Ferreira et al., Figure S3

### Figure S4

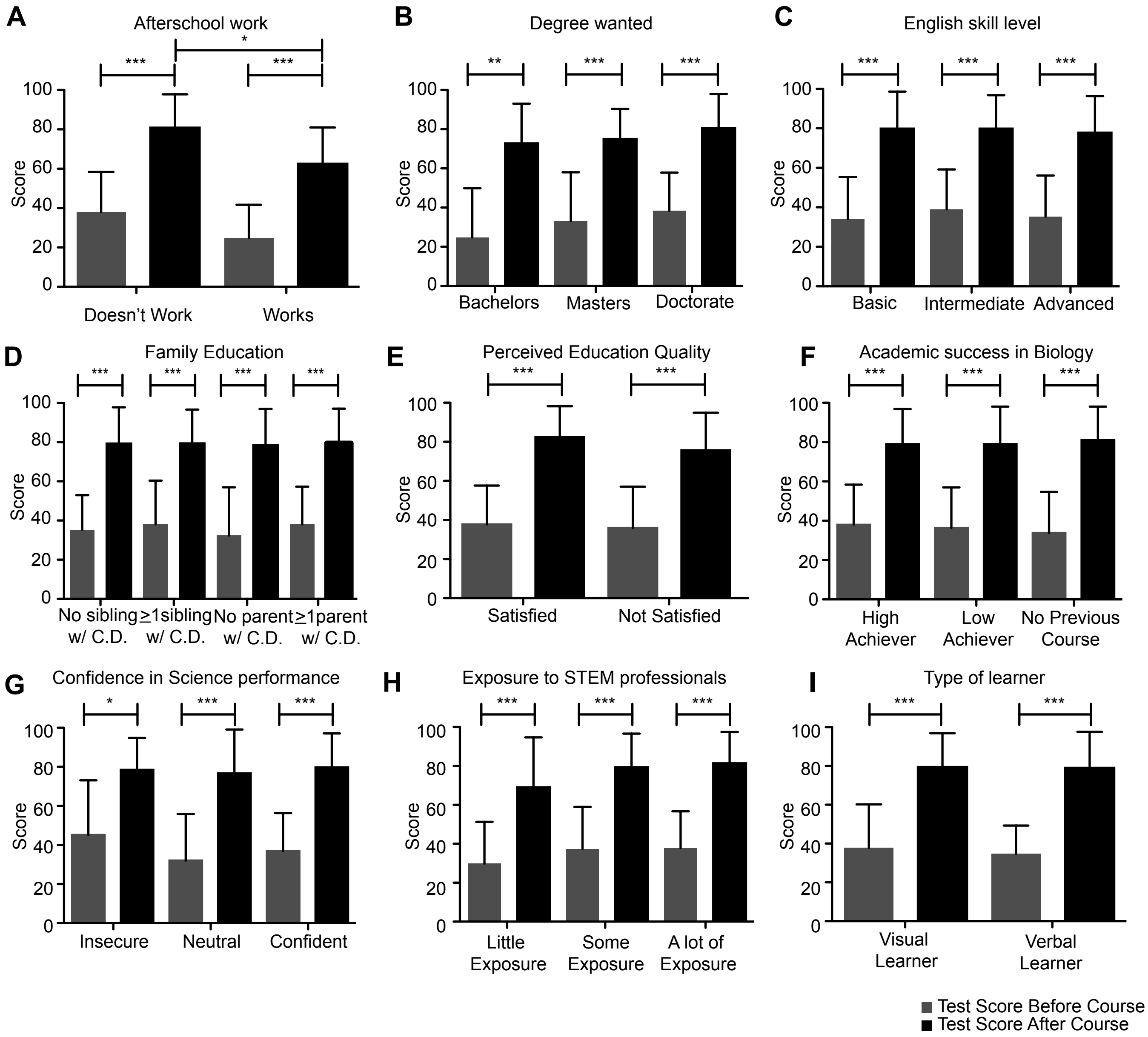


Ferreira et al., Figure S4
